# Ultrastructure of COPII vesicle formation characterised by correlative light and electron microscopy

**DOI:** 10.1101/2022.03.13.484130

**Authors:** Alejandro Melero, Jerome Boulanger, Wanda Kukulski, Elizabeth A. Miller

**Affiliations:** Cell Biology Division, MRC Laboratory of Molecular Biology, Cambridge, United Kingdom

## Abstract

Traffic of proteins out of the endoplasmic reticulum (ER) is driven by the COPII coat, a layered protein scaffold that mediates the capture of cargo proteins and the remodelling of the ER membrane into spherical vesicular carriers. Although the components of this machinery have been genetically defined, and the mechanisms of coat assembly extensively explored *in vitro*, understanding the physical mechanisms of membrane remodelling in cells remains a challenge. Here we use correlative light and electron microscopy (CLEM) to visualize the nanoscale ultrastructure of membrane remodelling at ER exit sites (ERES) in yeast cells. Using various COPII mutants, we have determined the broad contribution that each layer of the coat makes in membrane remodelling. Our data suggest that inner coat components define the radius of curvature whereas outer coat components facilitate membrane fission. The organization of the coat in conjunction with membrane biophysical properties determine the ultrastructure of vesicles and thus the efficiency of protein transport.

## Introduction

Eukaryotic cells use vesicular membrane carriers to shuttle proteins and lipids between subcellular compartments. These carriers are generated by cytosolic coat proteins capable of concentrating cargo at discrete membrane subdomains and enforcing membrane shape remodelling to generate highly curved vesicles and tubules (Bonifacino and Glick, 2004; Faini et al., 2013). Coat complex protein II (COPII) is responsible for the traffic of proteins from the endoplasmic reticulum (ER) to the Golgi complex, an essential route for 1/3 of the eukaryotic proteome. Vesicle formation occurs at ER exit sites (ERES), where the COPII machinery orchestrates cargo selection and membrane bending (Figure 1A) (Miller and Barlowe, 2010).

**Figure 1.**
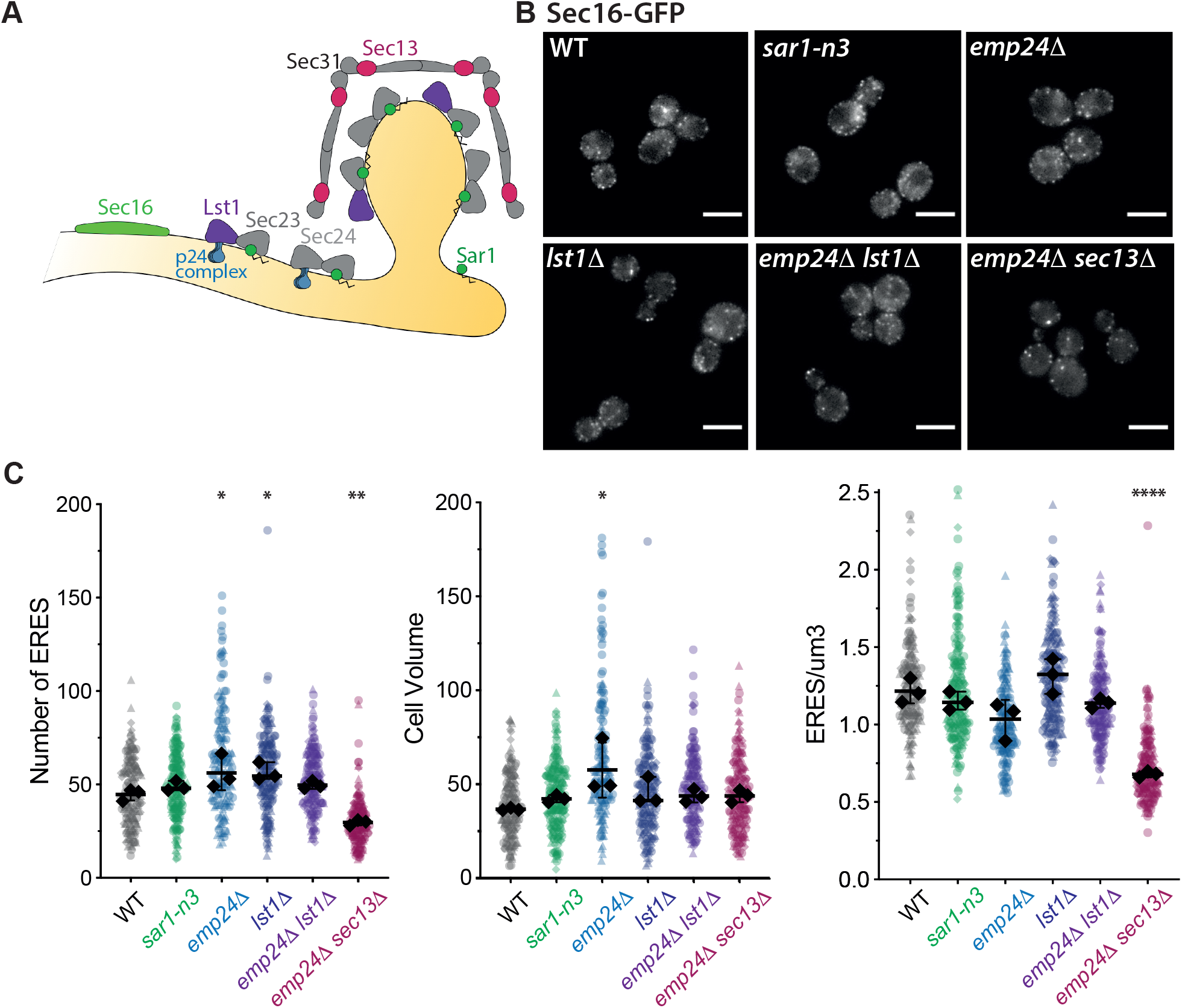
**A**) COPII assembly scheme. Sar1-GTP binds to the ER surface via an amphipathic helix. Sar1 recruits Sec23/Sec24, with Sec24 binding to cargo and cargo receptors (e.g. p24 complex). Sec23 also forms dimers with Lst1, a paralog of Sec24. Sar1-Sec23/Sec24 recruits Sec13/Sec31, which organizes as a rigid polyhedral structure. Sec16 is a regulatory protein which localizes at ERES and regulates coat polymerization. **B**) Visualization of ERES in the indicated yeast strains expressing Sec16-sfGFP. Scale bar is 5 μm. **C**) Superplots of quantification of ERES abundance and cell volume in COPII-dysfunctional strains, showing total number of ERES per cell (left panel), cell volume (μm^3^; middle panel), and ERS abundance per cell volume (ERES/μm^3^; right panel). Values for individual measurements are represented as dots, with dot shape indicating which of three biological replicas the sample came from. Black dots represent median values for each biological replica; bars indicate the mean and standard deviation between replicas; statistical significance was analyzed using a one-way ANOVA (* p<0.05; ** p=0.008; **** p<0.0001).

Vesicle formation starts with the small GTPase, Sar1, which upon activation inserts an amphipathic a-helix into the cytosolic leaflet of the ER membrane, enabling the recruitment of subsequent coat components. The inner coat is composed of the heterodimer Sec23/Sec24, where Sec23 is GTPase activating protein (GAP) for Sar1, and Sec24 is a cargo adaptor which interacts directly with cargo or with cargo receptors through their sorting signals. Together, Sar1-Sec23/Sec24 trigger the recruitment of the outer coat, composed of the rod-like heterotetrameric Sec13/Sec31. Assembly of the outer coat into a polyhedral cage is thought to drive membrane deformation into highly curved buds (Stagg et al., 2006), leading to the appearance of a narrow bud neck which undergoes fission, probably facilitated by rounds of Sar1 GTP hydrolysis (Bi et al., 2007; Hanna et al., 2016; Kurokawa et al., 2016).

Interactions between the outer and inner coat layers drive coat assembly, but also play a regulatory role in coat assembly. An intrinsically disordered region of Sec31 interacts with Sec23 via multiple weak interfaces. One interface accelerates GTP hydrolysis by promoting optimal amino acid positioning within the catalytic pocket (Bi et al., 2007). Additional roles may include coordinating the lateral assembly of the inner coat and stabilizing the coat scaffold (Hutchings et al., 2018; Stancheva et al., 2020). In addition to the core COPII coat proteins, various accessory proteins are thought to influence COPII assembly. For example, the essential protein Sec16, also contains intrinsically disordered regions, and modulates the GAP activity of the assembled coat by competing with Sec31 for binding to Sec23 (Kung et al., 2011). This activity is thought to stabilize coat assembly, which might be particularly important to prevent early scission and thus facilitate the formation of large vesicles.

The question of how vesicles of different sizes are generated by the COPII coat remains an important question. Multiple factors likely contribute, including the rate of GTP hydrolysis, which determines coat lifetime on the membrane (Antonny et al., 2001), and organization of the inner coat by accessory proteins (Ma and Goldberg, 2016; Raote et al., 2018). Moreover, distinct properties of the inner coat proteins also seem to influence vesicle size; in yeast, the Sec24 paralog, Lst1, increases the average size of COPII vesicles by ~15% (Gomez-Navarro et al., 2020; Miller et al., 2002; Shimoni et al., 2000). The mechanism by which Lst1 drives the formation of larger vesicles remains unknown. Tuning vesicle size to the appropriate cargo burden is important not just in accommodating large cargo proteins (Malhotra et al., 2015). We recently showed that the sorting capacity of COPII vesicles stems at least in part from steric effects: vesicles crowded with cargo proteins preclude the passive leakage of ER resident proteins (Gomez-Navarro et al., 2020). Thus, tuning the ultrastructure of vesicles to accommodate the requisite cargo and maximising vesicle occupancy is a fundamental aspect of protein quality control.

Despite our knowledge of the parts list of the coat machinery, and a growing understanding of the assembly process, precisely how the COPII coat overcomes the energy barrier to remodel a relatively flat and cargo-crowded membrane into a vesicle with distinct morphology remains unclear. The coat must overcome both the physical opposition conferred by cargo crowding in the lumen and the resistance of the lipid bilayer to changes in shape (Derganc et al., 2013; Stachowiak et al., 2013). *In vitro* evidence suggests that membrane remodelling can be driven by all components of the coat via different mechanisms (Hutchings et al., 2018; Lee et al., 2005; Zanetti et al., 2013). However, *in vitro* techniques fail to recapitulate the complexity of ER architecture and luminal crowding, hence limiting our understanding of the mechanistic requirements of COPII machinery. In cells, vesicle formation at ERES is very dynamic and vesicles are too small to be visualized by light microscopy. Traditional electron microscopy cannot reveal protein localization, and thus offers limited information on the ultrastructure of specific budding events. Here we investigate membrane ultrastructure during COPII vesicle formation using correlative light and electron microscopy (CLEM). We obtained insight into COPII vesicle formation by examining yeast mutants with compromised coat subunits. We find that different COPII coat subunits have specific roles in curvature and force generation, which ultimately determine ultrastructure and thus influence secretion. Our data support a model for COPII vesicle formation where the inner coat components define the radius of curvature, and the outer coat drives membrane remodelling to high curvature and fission.

## Results

### Inner and outer coat dysfunction modifies the ultrastructure of ERES

To understand the role of distinct COPII subunits in membrane remodelling at ERES we used yeast mutants defective in coat assembly, cargo packing or membrane remodelling (Figure 1A). These mutations do not compromise cell viability *in vivo*, suggesting that cells can adapt to limitations in coat function. To challenge early coat assembly and membrane remodelling we used a mutant Sar1 strain, *sar1-n3* (Lee et al., 2005). This mutation substitutes two bulky residues for alanine within the N-terminal amphipathic helix, which abolishes the capacity of Sar1 to deform membranes *in vitro* and reduces export of GPI-APs *in vivo* (Lee et al., 2005). We addressed inner COPII coat function with three different strains: *emp24*Δ, *lst1*Δ and *emp24*Δ *lst1*Δ. Emp24 is a member of the p24 family, which forms a cargo-receptor complex required for sorting of GPI-APs into COPII buds (Manzano-Lopez et al., 2014). We used an *emp24*Δ strain to reduce the abundance of bulky cargo at ERES and thus reduce the energy barrier to deform the ER membrane (Copic et al., 2012). GPI-APs bound to the p24 complex are preferentially selected for secretion by the cargo adaptor subunit Lst1, an isoform of Sec24 (D’Arcangelo et al., 2015; Manzano-Lopez et al., 2014). Because Lst1 is associated with large COPII vesicles (Miller et al., 2002; Shimoni et al., 2000), we used the *lst1*Δ background to force the formation of small buds and vesicles. The *emp24*Δ *lst1*Δ double mutant should yield small vesicles with reduced load of bulky cargo (Gomez-Navarro et al., 2020). Finally, we assessed the contribution of a rigid outer coat scaffold, by analysing a *emp24*Δ *sec13*Δ strain, where deletion of Sec13 is tolerated, likely because of reduced GPI-AP traffic, which lowers the energetic cost of bending the ER membrane (Copic et al., 2012).

To visualize ERES we tagged Sec16 at its chromosomal locus by integrating superfolder GFP (sfGFP) at the C-terminus (Sec16-sfGFP) (Figure 1B). We first asked whether the number and/or distribution of ERES was altered in any of our mutant backgrounds. We acquired image stacks and counted the number of ERES per cell (Figure 1C; left panel) and cell volume (Figure 1C; middle panel) using a custom-made FIJI macro. Only the *emp24*Δ *sec13*Δ strain showed significant alterations of the number of ERES per cell volume, with an abundance of ERES/μm^3^ ~40% lower than wild type (Figure 1C; right panel). Having a reduced capacity to deform membranes (*sar1-n3*), or making smaller (*lst1*Δ) and/or less crowded vesicles (*emp24*Δ *lst1*Δ and *emp24*Δ) did not impact the number or distribution of ERES in yeast cells.

To gain a detailed understanding of membrane ultrastructure at ERES we used correlative light and electron microscopy (CLEM). This method combines the advantages of protein localization by light microscopy with the detailed nanoscale resolution of electron tomography (Kukulski et al., 2011). We have shown recently that CLEM is a suitable tool to analyse the ultrastructure of COPII vesicles (Gomez-Navarro et al., 2020). Yeast cells were subjected to high pressure freezing, freeze substitution and embedding in Lowicryl resin. This approach enables excellent preservation of ultrastructural details, while allowing the use of room temperature electron microscopy to localize a protein of interest within a tomographic volume (2×2 μm by ~300 nm in depth). We acquired fluorescence microscopy images of cells in resin sections, and used fluorescent fiducials to obtain a high precision correlation of ERES marked with Sec16-GFP within electron tomograms (Figure 2A). Using this approach, we analysed the ultrastructure of ERES in wild-type and mutant cells (Figure 2B).

**Figure 2.**
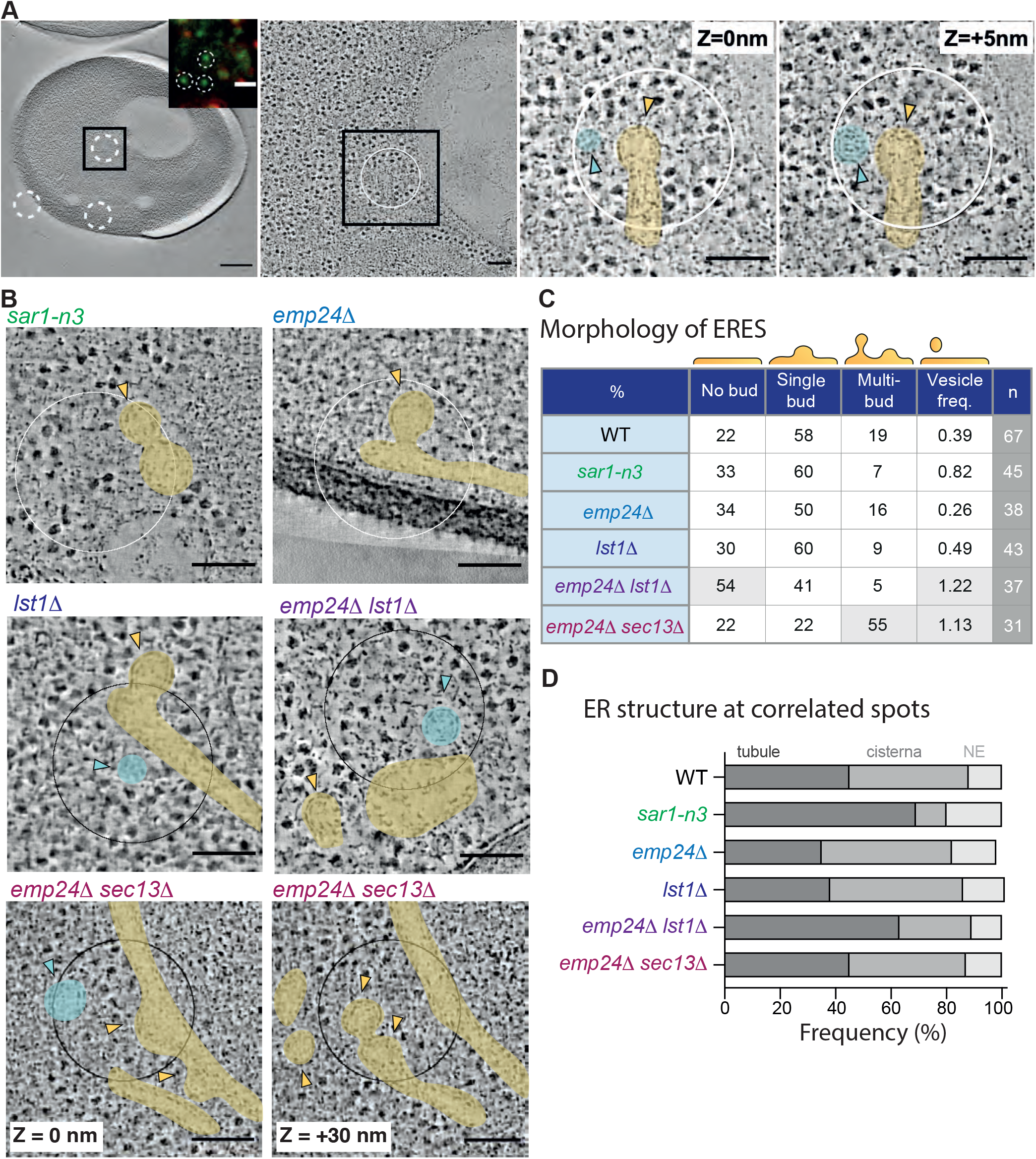
**A**) CLEM workflow: sections of high-pressure frozen and resin-embedded yeast cells were imaged with fluorescent microscopy (inset, left panel) and scanning transmission electron microscopy (STEM). Fluorescence images were correlated with low-magnification EM images (left panel) using fluorescent fiducials (red spots in inset), enabling the precise localization of Sec16-sGFP positive ERES (green spots in inset; white circles in EM images) within high-magnification tomographic volumes (right panels). Vesicle formation from ER structures (yellow arrow) and free vesicles (cyan arrow) can be found at sites of Sec16-sfGFP signal. **B**) Examples of correlated Sec16-sfGFP ERES in COPII dysfunctional strains. For *emp24*Δ *sec13*Δ we show two virtual slices of the same tomogram. **C**) Table of ERES ultrastructure categories as percentage of number of correlated Sec16-sfGFP (n) subcellular regions; vesicle frequency represents the average number of free vesicles per ERES, quantified as the total no. of free vesicles divided by the total number of ERES. Grey shaded sections represent major differences from WT. **D**) Bar plot representing the frequency (%) of Sec16-correlated spots in different ER sub-regions. In the absence of visible buds, the nearest ER membrane to the centroid of the Sec16-sfGFP signal was considered as the corresponding ER sub-region. Scalebars in A: 500 nm in left panel (1μm in inset) and 100 nm in subsequent panels. Scale bars in B: 100 nm.

Tomographic reconstructions are detail-rich, enabling us to extract varied information about the environment of ERES as well as their ultrastructure (Figure 2C, 2D). The majority of wildtype ERES (~60%) comprise a single bud; occasionally, multiple budding events could be found at the same Sec16-sfGFP correlated region (19%), although we did not observe more than 2 simultaneous buds per ERES (Figure 2C). Free vesicles fully released from the donor membrane could be observed, with a frequency (0.39 for WT) that we define as the number of vesicles per ERES (i.e. the total number of free vesicles divided by the number of correlated ERES). Sec16-correlated regions occurred with equivalent frequency on ER tubules or cisterna, often at regions of existing curvature (fenestrations or membrane junctions), with a minority of ERES found at the nuclear envelope (NE) (Figure 2D). We also note the presence of other nearby structures, such as Golgi cisterna (Supplementary Figure 1).

Comparing the global features of ERES in different mutant backgrounds (Figure 2C, 2D), we highlight several significant differences. Mutation of the Sar1 N-terminal amphipathic helix (*sar1-n3*) reduced the occurrence of multi-budded structures (to 7%), and increased the frequency of free vesicles (~0.8) associated with ERES (Figure 2C). Budding sites in this background were more prevalent on ER tubules (Figure 2D), suggesting ERES form preferentially on existing areas of curvature when membrane remodelling is compromised (Okamoto et al., 2012).

The *emp24*Δ strain was broadly similar to wild-type with respect to donor membrane morphology and distribution, whereas the *lst1*Δ strain had reduced occurrence of multi-budded structures (Figure 2C). In contrast, the *emp24*Δ *lst1*Δ double mutant markedly differed from both wild-type and individual deletions. The frequency of free vesicles (1.22) was 3 times higher than wild type, corresponding to an average of more than 1 free vesicle per ERES, and the predominant budding profile was a flat membrane (Figure 2C), suggesting increased efficiency or kinetics of vesicle release in the double mutant. Budding sites in the *emp24*Δ *lst1*Δ double mutant were more frequently found on tubular ER (Figure 2D). Finally, the budding profile of the *emp24*Δ *sec13*Δ strain, with a compromised outer coat, was shifted towards bud production, with 55% of ERES occurring as multiple buds (Figure 2C). In some cells, these structures formed a complex network of tubules decorated with buds and pearled-tubes (Supplemental Figure S1). ERES in *emp24*Δ *sec13*Δ cells were also associated with significantly more free vesicles per ERES (vesicle frequency of 1.13, corresponding to an average of 1 vesicle per ERES). Our CLEM tomography analysis of ERES thus revealed that the ultrastructure of vesicle biogenesis adapts to the physical features of the coat and the cargo load. Altering inner coat function (*sar1-n3* and *emp24*Δ *lst1*Δ stimulates vesicle release resulting in reduced steady-state curvature at ERES, whereas reducing the rigidity of the outer coat prolongs bud formation resulting in extended multi-budded structures.

### Ultrastructural changes in COPII vesicles suggest distinct roles in membrane remodelling and scission for inner and outer coat layers

We next sought to quantify vesicle morphology by obtaining quantitative morphological descriptors. We first used a semi-automated segmentation method to segment the membrane boundaries of free vesicles in tomographic reconstructions (Figure 3A; (Machado et al., 2019)). Using the segmented surface mesh, we calculated the volume of the vesicle (Figure 3B), determined the centroid of the vesicle, and measured the furthest and closest distance from the centroid to the vesicle surface, equivalent to the major and minor radii of the vesicle (Figure 3A). As reported previously (Gomez-Navarro et al., 2020), vesicle volume was reduced in *emp24*Δ *lst1*Δ mutants, and increased in *emp24*Δ *sec13*Δ double mutants (Figure. 3B). As expected, we saw similar reduction in vesicle volume in the *lst1*Δ single mutant, with volumes unchanged in *sar1-n3* and *emp24*Δ mutants (Figure 3B).

**Figure 3.**
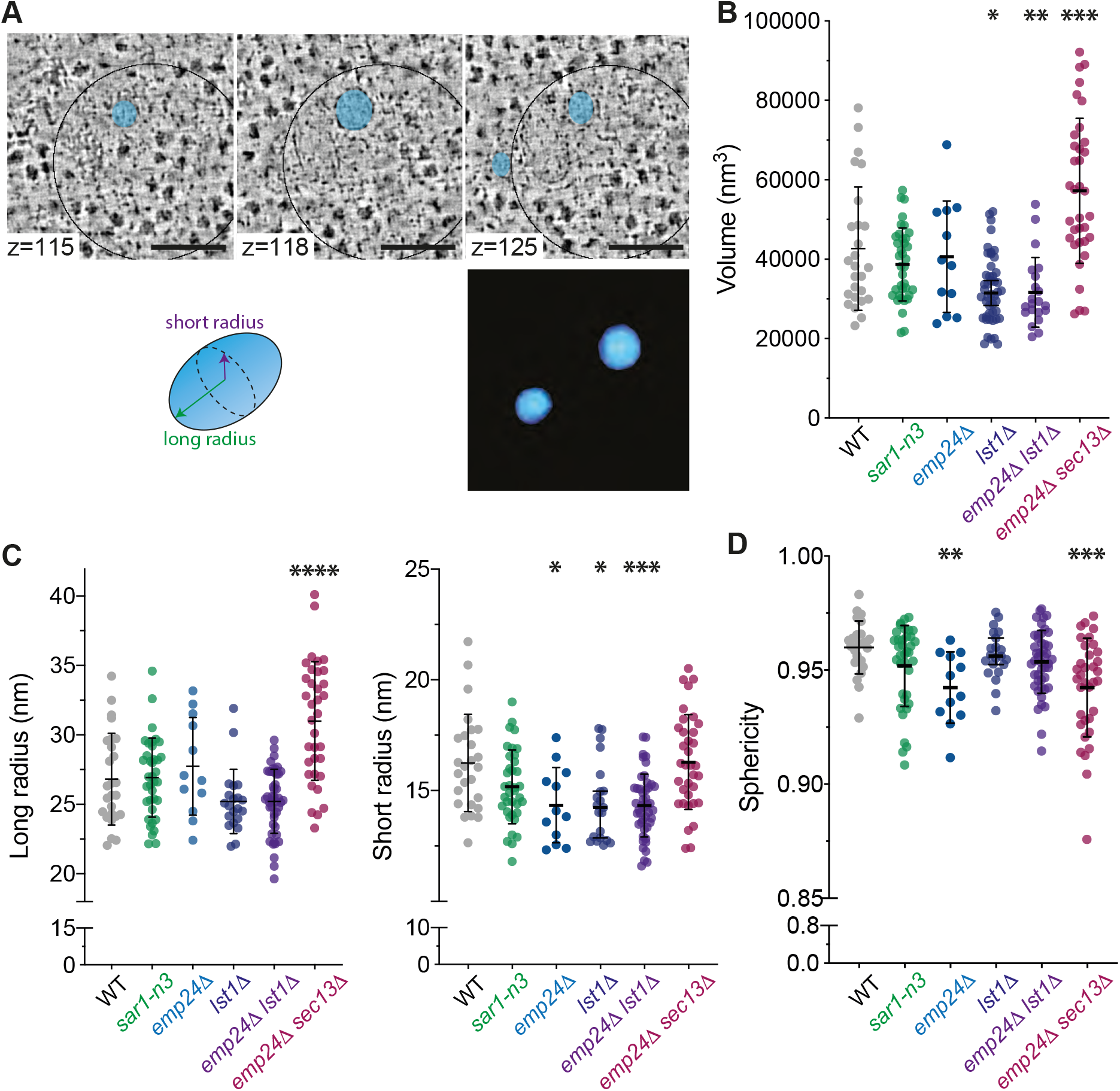
**A**) Upper panels: Sequential virtual slices of a Sec16-sfGFP correlated site showing two free COPII vesicles (false coloured in cyan) next to a budding site. Scalebars are 100 nm. Lower left; diagram defining axes of ellipsoidal vesicles. The centroid of the vesicle was used to calculate the longest radius (farthest point to the centroid) and the shortest radius (nearest point to the centroid). Lower right; segmented volume of upper panel vesicles. **B**) Plot of the volume (nm^3^) of free COPII vesicles in the strains indicated. Each point corresponds to a vesicle measured in the indicated strain. Bars represent median value and 95% confidence interval. **C**) Analysis of the two major axes of free COPII vesicles. Using the segmented volumes of COPII vesicles we determined the longest and shortest radius from the centroid for each vesicle in wild type and COPII-dysfunctional strains. Each point corresponds to a vesicle analyzed in the indicated strain. Bars represent median with 95% confidence interval. **D**) Plot of the sphericity of vesicles, where 1 equals a perfect sphere and <1 indicates ellipsoidal morphologies. Statistical tests were one-way ANOVA with Tukey’s correction for multiple comparisons (* p<0.05; ** p<0.01; ** p<0.001; **** p<.0001).

Examining the ellipsoidal nature of free vesicles, we found that wild type vesicles range from ~25-35 nm in their major radius, with the minor radius lying between ~15-25 nm (Figure 3C), making COPII vesicles ~1.5 times longer than wider on average. Measuring the vesicle dimensions in COPII mutants using this approach revealed that the *emp24*Δ *sec13*Δ strain accumulates vesicles ~10% larger than wild type cells, with the difference driven by an increase in the major axis (Figure 3C). On the other hand, we found that almost all inner coat mutants (*lst1*Δ, *emp24*Δ *lst1*Δ, *emp24*Δ) had a significantly smaller minor radius relative to wild type vesicles, with a median of ~15 nm. Vesicles in these strains had no significant difference in major radius compared to wild type, although the range of length in *lst1*Δ strains was smaller than that of wild type cells (Figure 3C). The sphericity of free vesicles was calculated from the major- and minor-axis measurements, where a true sphere has sphericity of 1. Wild type vesicles are modestly non-spherical, with mean sphericity of 0.96 (Figure 3D); vesicles in *lst1*Δ, *emp24*Δ *lst1*Δ and *sar1-n3* strains showed similar sphericity (~0.95). In contrast, vesicles in *emp24*Δ and *emp24*Δ *sec13*Δ strains were significantly less spherical, reflected by their longer major axis relative to the minor axis (Figure 3C, D).

Together, our measurements of vesicles that lack Lst1 suggest that vesicles made with Sec24 as the dominant inner coat subunit have reduced vesicle volume due to tighter curvature along one radius. Exactly what features of Lst1 and Sec24 drive these differences in membrane curvature generation remain to be tested. Our data also suggest that the outer coat subunit, Sec13, predominantly drives vesicle fission; in the absence of Sec13, vesicle length increases with no change in the minor radius, consistent with a delay in vesicle release. The analysis of *emp24*Δ vesicles is intriguing; despite having all coat components, these vesicles are less spherical (i.e. flatter or more ellipsoidal) than wild type vesicles. Flatter vesicles suggests that the low abundance of bulky cargo indeed facilitates membrane bending, perhaps due to low inner pressure within the vesicle lumen (Gomez-Navarro et al., 2020). Finally, our analysis shows that *sar1-n3* vesicles are not morphologically different to those of wild type cells, despite *in vitro* evidence that shows significant defects in membrane remodelling (Lee et al., 2005). One explanation for this apparent discrepancy is that ERES in the *sar1-n3* mutant are more abundant on tubular ER (Figure 2D), which has significant intrinsic curvature. Tubular ER has an average radius similar to that of intermediate and late budding events (~20 nm; discussed further below) (West et al., 2011), suggesting that the COPII coat can remodel tubes with less energy cost than on flat cisternae (Okamoto et al., 2012). Together, these measurements suggest that Sar1 may be involved in events other than curvature generation, which seems to be more dominantly associated with other coat components.

### Quantitative analysis of early stages of vesicle formation

Our findings suggest that inner and outer coat play distinct roles in membrane remodelling. We next measured the ultrastructure of budding intermediates attached to the ER membrane in order to obtain quantitative information about membrane remodelling during the budding process. Buds are open contours that cannot be segmented with the approach used for free vesicles. We therefore used an alternative approach where we manually measured bud length and radius of curvature. The radius of curvature was extracted by fitting a circle to the tomographic plane with the largest bud diameter and determining the radius of that circle, whereas bud length was calculated by measuring the straight-line distance between bud tip and the ER baseline (Figure 4A). We propose that these measurements broadly recapitulate bud progression, with large radius of curvature and low bud length early during vesicle formation, followed by a decrease in bud radius and increase in bud length (Figure 4B). Looking first at the global measurements of bud length and radius, we find that in wild type cells, the average bud is 40 nm long with a radius of 28.5 nm (Figure 4C, D). The *emp24*Δ and *emp24*Δ *sec13*Δ strains had broader distributions of bud lengths than wild-type cells, with the mean bud length significantly larger (Figure 4C). Interestingly, *lst1*Δ suppressed the effect of *emp24*Δ on bud length, with the *emp24*Δ *lst1*Δ strain resembling wild type (Figure 4C). Other inner coat mutants (*lst1Δ*, and *sar1-n3* strains) also had bud lengths similar to wild type (Figure 4C). The average bud radius was similar across all strains (Figure 4D).

**Figure 4.**
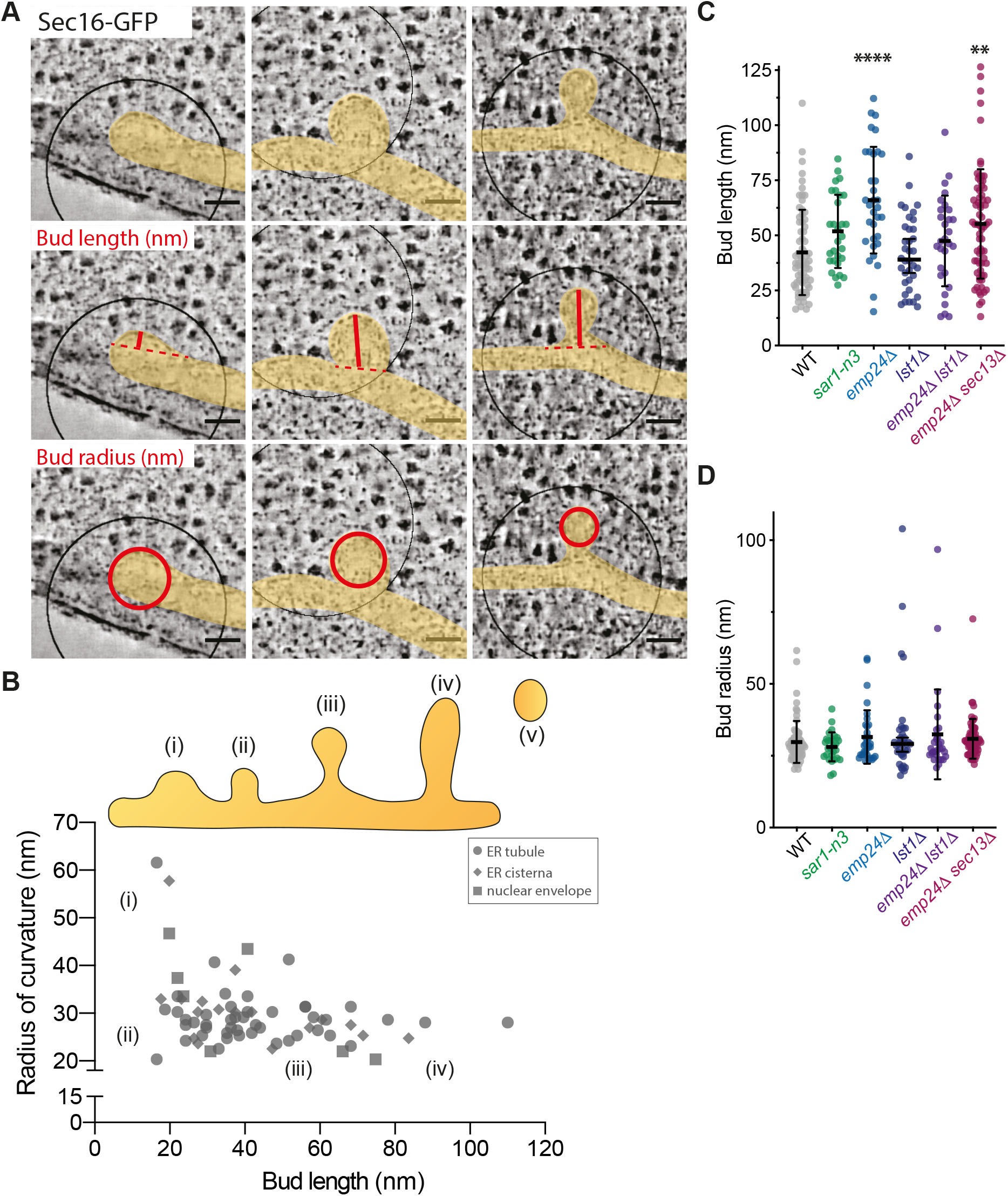
**A**) Measurement of bud ultrastructure at WT ERES (black circle represents a 250 nm circle around the centroid of the Sec16-GFP spot; ER membranes false coloured in yellow). Bud length and bud radius were calculated from tomograms of various budding morphologies including shallow buds (left panels), cup-shaped structures (middle panels) and omega-shaped structures/tubes (right panels). Bud length corresponds to the distance between the tip of the bud to the ER baseline (solid red lines), whereas bud radius was detemined by fitting a circle at the equatorial plane of the budding event (solid red circles). **B**) Distribution of WT budding events as a function of bud length and radius. Bud maturation steps can be categorized as (i) shallow buds of low curvature and short length, (ii) early buds of high curvature and short length, (iii) late buds with high curvature and elongated bud-neck, and (iv) very long buds with high curvature. Each point represents a single budding event in a wild-type strain with the ER morphology at the bud site indicated by the shape of the datapoint. **C**) Plot of bud length (nm) of individual buds in the indicated strains. Bars represent the median with 95% confidence interval. **D**) Plot of bud radius (nm) of individual buds in the indicated strains. Bars represent the median with 95% confidence interval. Statistical test was a oneway ANOVA with Tukey’s correction for multiple comparisons (* p<0.05; ** p<0.01; ** p<0.001; **** p<.0001).

By plotting bud length against radius of curvature for each individual bud, we obtained the budding profile of COPII in wild type ERES (Figure 4B). Early buds are ribosome-free patches of ER with high radius of curvature and low bud length. This stage is likely followed by an increase in bud length and curvature, with buds maturing into cup-shaped structures of 20 to 30 nm radius and ≤40 nm in length. The increase in curvature is followed by an increase in length up to 80 nm including bud-neck. Knowing the average vesicle length (Figure 3), we can infer that fission takes place 50-60 nm from the bud tip. We classified each bud according to the morphology of the donor ER membrane, finding that the tubular, cisternal or nuclear envelope origin of the membrane did not seem to correlate with a particular budding morphology (Figure 4B).

This simple yet detail-rich approach enabled us to compare the maturation of COPII buds between wild type and coat mutant strains (Figure 5). Looking first at the *sar1-n3* strain, we find the absence of both shallow curvature stages and very short bud lengths (Figure 5A). We note that the majority of these budding events took place at curvature-rich tubular ER (Figure 5F); bud initiation on already curved membranes with ready progression to higher curvature could explain the low frequency of shallow buds. In the *emp24*Δ strain, in which bulky cargo are decreased in vesicles, we see few buds with high curvature and short bud length, but more long highly curved buds than in wild type. (Figure 5B). Since bud length is on average longer in this strain (Figure 4C), one interpretation is that later stages of bud formation and/or vesicle fission represent a bottleneck such that these late-stage intermediates accumulate more than in wild-type cells. In *lst1*Δ and *emp24*Δ *lst1*Δ strains, where Sec24 is the dominant inner coat subunit, membrane remodelling seems to follow a profile similar to wild type, perhaps with a more acute transition from shallow to high curvature (Figure 5C,D). Finally, the *emp24*Δ *sec13*Δ strain showed reduced frequency of shallow-curved buds, with most budding events apparently in mid/late stages (Figure 5E). As described earlier (Figure 2C), many of these buds were found in ERES with multiple budding sites, which suggests membrane remodelling and curvature generation is not a limiting step in the absence of Sec13.

**Figure 5.**
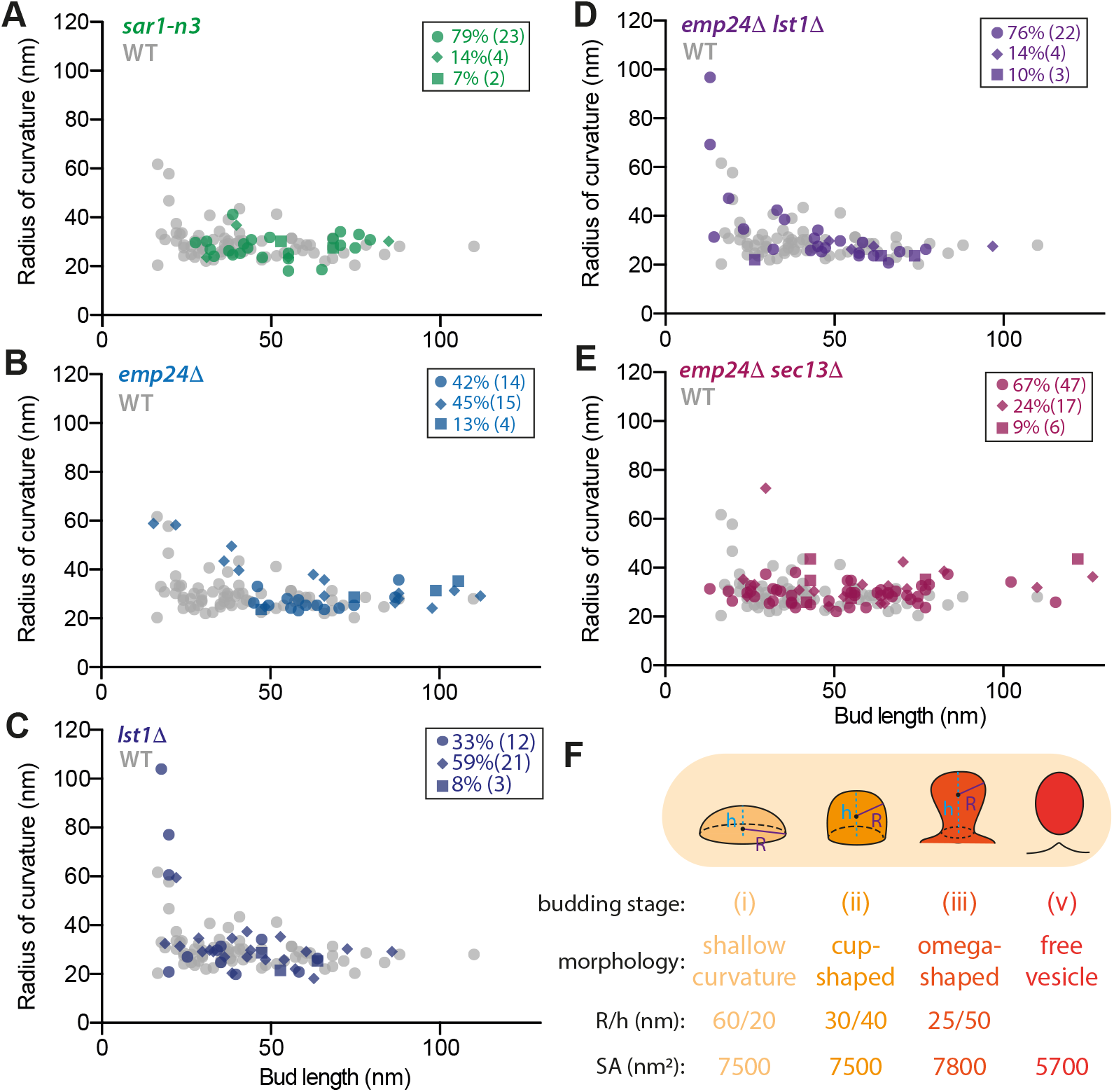
**A - E**, Bud morphology as described for Figure 5B comparing wild-type (grey datapoints) and indicated strains (coloured datapoints) showing the distribution of budding events as a function of bud length (nm) and radius of curvature (nm). Inset panels indicate the bud origin, where circle = ER tubule, diamond = ER cisterna, and square = nuclear envelope. **F**) Bar plot indicating the frequency (%) of buds as per ER sub-region in wild-type and dysfunctional COPII strains.

Our analysis of budding intermediates still attached to the donor membrane suggests that ER remodelling progresses with a relatively constant surface area (Figure 5F), such that the radius of curvature and bud length simultaneously increase over time. Early-stage intermediates with shallow curvature (60 nm radius, 20 nm height) have a curved surface area (~7500 nm^2^), which likely corresponds to a COPII-coated region, that closely matches that of buds with cupshaped structures with no obvious bud-neck (~30 nm radius, 40 nm height; surface area ~7500 nm^2^). Longer buds with a characteristic omega shape have a smaller radius of curvature and a narrow bud-neck (~25 nm radius and ~50 nm in length, including bud-neck) yielding a surface area of ~7800 nm^2^. The extra surface area of this structure likely includes uncoated regions such as the bud neck. Free vesicles have an average surface of ~5700 nm^2^ (Gomez-Navarro et al., 2020), suggesting some loss of membrane surface area is associated with vesicle fission and release. Electron cryo-tomography will be required to directly visualize coats on these curved membranes.

## Discussion

CLEM is a powerful tool to investigate the ultrastructure of membranes within cells. Here, we have measured various properties of vesicles and budding intermediates at ERES in yeast. Genetic modification of coat components or cargo abundance significantly modifies the behaviour of the coat and subsequently the ultrastructure of buds and vesicles. By systematically analysing membrane features in various COPII mutants, we have obtained new insight into the contributions of different components.

A key finding of our CLEM analysis of coat mutants is that the inner coat likely plays a significant role in membrane remodelling. Pioneering *in vitro* reconstitutions and structural studies have long suggested that the mechanical force to bend the ER membrane comes predominantly from the rigid cage of Sec13/Sec31 (Bacia et al., 2011; Barlowe et al., 1994). More recent structural evidence, however, indicates that assembly of the inner coat (Sar1-Sec23/Sec24) might also drive membrane remodelling, whereby organization into helical arrays is associated with tubulation of large synthetic liposomes (Hutchings et al., 2021; Zanetti et al., 2013). Consistent with a significant role for the inner coat in curvature generation, our tomography revealed that lack of Sec13, which likely results in a flexible Sec31 cage, does not prevent the formation of buds and vesicles with high curvature. The caveat to this conclusion is that loss of bulky cargo (i.e. *emp24*Δ) is required for this permissive membrane bending condition.

If a rigid Sec13/31 cage is not absolutely required to achieve high curvature, what is the role of the outer coat in membrane remodelling and why does Sec31 stiffness matter? Our analysis of bud and vesicle length might shed light on this. Although both *emp24*Δ and *emp24*Δ *sec13*Δ mutants have long buds, only in the *emp24*Δ *sec13*Δ does this result in elongated vesicles, which suggests that the absence of Sec13 delays fission. Consistent with such a model, a majority of *emp24*Δ *sec13*Δ ERES are multi-budded structures, suggesting defects in vesicle release. We propose that membrane remodelling towards shallow curvature is driven by inner coat assembly; a rigid Sec13/Sec31 scaffold then encapsulates the membrane to form a structure with an elongated bud-neck, which would facilitate spontaneous or Sar1-mediated fission (Figure 6). A less rigid cage would lack the necessary structural stiffness to counteract pressure within the bud, thereby generating larger buds without a narrow bud-neck where fission could occur.

**Figure 6.**
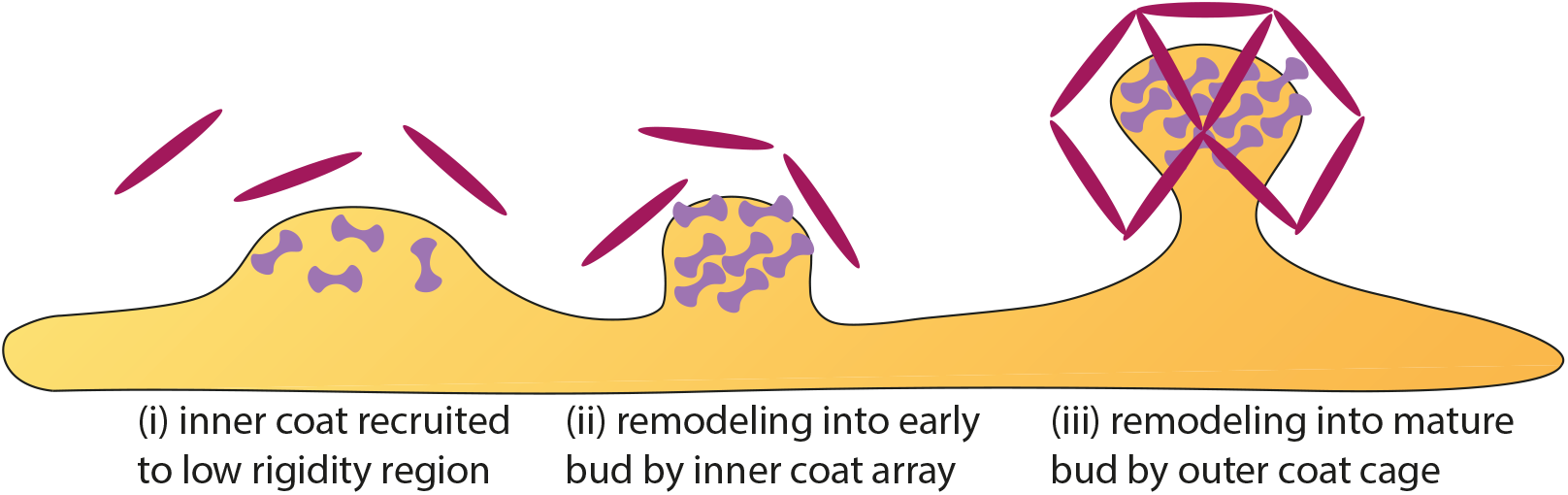

### Model

The capacity of the coat to sense and modify the physical properties of the membrane seems to be critical to both initiate a budding site and determine vesicle ultrastructure. Neither bending rigidity nor membrane tension are uniform, either throughout the cell or over the remodelling process. Thus, the coat likely needs diverse strategies to impose the required shape changes needed to ultimately drive fission. Low membrane bending rigidity and high spontaneous curvature are desirable features to initiate budding. ER ultrastructure is actively remodelled via shape changes and insertion of new lipids, both of which may favour the spontaneous appearance of conditions suitable for budding. Sar1 might sense such favourable conditions (Doucet et al., 2015) (intrinsic curvature and low bending rigidity) and together with Sec16 drive COPII recruitment to suitable regions of the ER. In yeast, the ER membrane is studded with numerous ribosomes, which need to be cleared to permit coat assembly and membrane remodelling. How ribosome clearance is coordinated with COPII recruitment remains unclear, but the large size and intrinsic disorder of Sec16 suggest one of its roles may be to prepare an appropriately cleared region of ER membrane.

Once bud site selection has occurred, local recruitment of the inner coat would initiate membrane bending, driven by lateral assembly of Sar1-Sec23-Sec24 units. As the bud reaches high curvature, the steric pressure associated with locally enriched adaptor-bound cargo crowding increases the energetic cost to change membrane curvature (Derganc et al., 2013). Sar1 may locally reduce the bending modulus of the lipid bilayer and facilitate spontaneous positive curvature (Hanna et al., 2016; Lee et al., 2005; Loftus et al., 2012; Settles et al., 2010) thus counter-balancing the growing repulsive forces within the ER lumen as the coat polymerizes (Copic et al., 2012). Moreover, the lipid environment may be locally remodelled with lipid species that facilitate membrane bending or stabilize cargo within the unfavourable conditions of vesicle lumen (Jiménez-Rojo et al., 2019; Melero et al., 2018; Rodriguez-Gallardo et al., 2020). Recruitment of Sec13/Sec31 rods would promote the ordered arrangement of the inner coat (Hutchings et al., 2021; Ma and Goldberg, 2016), thus reducing the bud radius (Gomez-Navarro et al., 2020). Polymerization of Sec13/Sec31 into a rigid cage around the bud tip would further increase the pressure within the bud, which may translate into a local increase in surface tension and appearance of a bud-neck (Raote et al., 2020). The bud-neck is probably free of coat and cargo, but will be mechanically stressed, thereby facilitating spontaneous fission of the membrane and release of a free vesicle. This model explains our observations of high frequency of vesicles in *lst1*Δ and *sar1-n3* strains, where small bud radius (*lst1*Δ) and limited capacity to reduce rigidity (*sar1-n3*) may promote early membrane fission and vesicle release. Our model is also in agreement with previous *in vitro* observations, where COPII has been found to favour polymerization on liposomes with low membrane bending rigidity (Melero et al., 2018), similar to other coat complexes (Manneville et al., 2008; Mercier et al., 2020; Saleem et al., 2015).

Given the conserved structures of COPII coat subunits across species, our observations in yeast are likely relevant to other organisms, despite clear differences in the morphology of ERES. In animal cells, secretion of very large cargo such as pro-collagen fibres is difficult to reconcile with classic COPII secretion models (McCaughey et al., 2018). A recent theoretical study suggests that modulating bud tension could prevent fission and lead to the formation of pearled buds, which would support export of pro-collagen fibres (Raote et al., 2020). Our data supports the preferred formation of pearled buds if the physical stiffness of the Sec13/Sec31 cage is reduced (Figure 2C, Suppl. Fig. 2 and (Gomez-Navarro et al., 2020)), thus supporting the possibility of such an export mechanism for large cargo in animal cells. Recent studies have described large tubular and multi-budded ERES in human cells, where COPI machinery may also be required for ER-to-Golgi trafficking (Shomron et al., 2021; Weigel et al., 2021). Weigel and colleagues found that a sudden accumulation of cargo increases the volume of ERES, which may hinder the capacity of the coat to induce membrane fission, similar to the situation we observe in *emp24*Δ *sec13*Δ cells. Further application of CLEM approaches, combined with genome editing to introduce specific COPII coat mutants will help illuminate whether the principles of coat assembly and membrane remodelling we propose here are conserved in human cells.

## Acknowledgments

We thank Felix Campelo and Xiaohan Li for many helpful discussions, and the LMB EM facility for technical and logistical support. This work was supported by the Medical Research Council, as part of United Kingdom Research and Innovation (also known as UK Research and Innovation) under awards MC_UP_1201/10 to EAM and MC_UP_1201/8 to WK. For the purpose of open access, the authors have applied a CC BY public copyright licence to any Author Accepted Manuscript version arising.

## Materials and methods

### Strains and plasmids

All strains were generated and maintained using standard *S. cerevisiae* methods. Strains (Table S1) were made by PCR-based integration of sfGFP fused in-frame at the 3’ end of the *SEC16* locus using drug-resistance markers. Strain *emp24*Δ *sec13*Δ *SEC16-sfGFP pSEC13-URA3* was plated onto media containing 5-FOA to counter select for the *SEC13-URA3* plasmid.

### GFP imaging

For imaging of Sec16-sfGFP, cells were grown at 25°C in minimal media lacking tryptophan. Images were taken on a Nikon Ti2 with a 100×/1.49 NA Oil (TIRF) objective and a Hamamatsu ORCA-FLASH 4.0 C11330-22C camera using a scientific complementary metal-oxide-semiconductor (sCMOS) sensor. The same imaging methodology was used for the imaging of EM grids with sections of resin-embedded cells.

### ERES analysis

Z-stacks using 0.2 μm z spacing of epifluorescence images were acquired as described above. Cells were manually selected with the elliptical selection tool in FIJI and assigned as a region of interest. ERES were then localized with a difference of Gaussian (DoG) using a custom FIJI script. The local maxima of the DoG and a threshold based on a false alarm rate were combined to localize bright local maximas. Localized spots were then rendered as Gaussian spots and added as a second channel for visual validation. Statistical analysis was performed with Prism 7.0 (GraphPad Software).

### CLEM

CLEM was performed as described in Kukulski et al. (2011) with some modifications described in Ader and Kukulski (2017). Yeast cells were grown at 25°C in minimal media lacking tryptophan to 0.6–0.8 OD600 and pelleted by vacuum filtration onto nitro-cellulose discs, then placed on an agar plate to prevent the pellet from drying out. The yeast paste was high-pressure frozen in 200-μm-deep wells of aluminum carriers (Wohlwend) using a HPM100 (Leica Microsystems). Freeze substitution and Lowicryl HM20 (Polysciences, Inc.) resin embedding were done as previously described in Kukulski et al. (2011), with minor modifications. 0.03% uranyl acetate in acetone was used for freeze substitution. Samples were shaken on dry ice for the first 2–3 h of freeze substitution (McDonald and Webb, 2011). Sections of 300-nm thickness were cut with an Ultra 45° diamond knife (Diatome) on an Ultracut E microtome (Reichert). The sections were floated onto PBS and picked up with 200 mesh carbon-coated copper grids (AGS160, Agar Scientific). Fluorescent TetraSpeck beads (Invitrogen), 50 nm in diameter, were adsorbed onto the grid. Immediately after sectioning, grids were mounted for fluorescence microscopy (described above). Prior to electron tomography, 15 nm gold beads (Electron Microscopy Sciences) were adsorbed onto the sections, which were then post-stained for 15 min with lead citrate. Scanning transmission EM tomography was done on a TF20 microscope (FEI) with an axial brightfield detector, using a camera length of 200 mm and a 50-mm C2 aperture (Ader and Kukulski, 2017; Hohmann-Marriott et al., 2009). For correlation to fluorescence images, low magnification tilt series at 3.1 nm pixel size were acquired using SerialEM (±55° tilt range, 2° increment, single axis acquisition; Mastronarde, 2005). Higher magnification tomograms were acquired as dual axis tilt series ±60° with 1° increment and at 1.1 nm pixel size (Mastronarde, 1997). All tomographic reconstructions were done in IMOD (Kremer et al., 1996), and fiducial-based correlation was done using MATLAB-based scripts described in Kukulski et al. (2011).

After correlation of the Sec16-sfGFP signal, a circular area of 125 nm radius was placed on the centroid of the correlation. Buds and vesicles within this circular area were considered as part of an ERES. Frequency of free vesicles in the correlated region was calculated as total number of vesicles divided by the total number of correlated sites. For ER morphology determination, ER was considered cisternal when two parallel ER membranes were observed across the z axis, fusing at a cisterna edge only once. Tubular shape was designated when non-nuclear envelope membranes were not cisternal, generally corresponding to a hollow quasi-cylindrical structure of various diameters following irregular shapes across the z axis of the tomogram.

### Segmentation analysis

Free vesicles (Fig. 3) were manually segmented using IMOD. Vesicles axis and volume quantifications were obtained with FIJI using the plugin LimeSeg (Machado et al., 2019) as follows: the outer contour of a vesicle was selected as the region of interest (ROI) using the “point tool” and “segmented line” tool moving in z through the tomographic slices. The lowest plane of the vesicle was annotated as a ROI with the point tool, then contours were selected with the segmented line tool every five virtual slices before finally closing the volume with the top plane of the vesicle using the point tool. LimeSeg Skeleton Segmentation tool settings were adjusted to recognize and segment the outer surface of the vesicle (D_0: 4, F_pressure: 0, Z_scale: 1, Range_in_DO_u-nits: 1, NumberOfIntegrationStep: −1, RealXYPixelSize: 1). After running the segmentation, the correct distribution of surfels over the outer contour of the vesicle was assessed by eye. The LimeSeg segmentation tool provides the list of vertices of the mesh. The centroid of this point cloud gives an estimate the center of the segmented vesicle. The maximum radius was then computed as the maximum distance to this center considering the 1.1 × 1.1 × 1.1 nm voxel size. Sphericity was determined according to the formula (π^1/3*(6V)^2/3)/A. Statistical analysis was performed with Prism 7.0 (GraphPad Software).

### Bud surface area estimation

The surface area of early and medium buds was calculated using the formula SA = 2πRh, where R is the bud radius and h is the bud length. The surface area of late buds was calculated assuming a nearly spherical ellipsoid, using the expression SA ≈ 4π ^1.6^√(a^1.6^b^1.6^ + a^1.6^c^1.6^ + b^1.6^c^1.6^)/3. All calculations were made using https://www.calculator.net/surface-area-calculator.html.

---

**Supplementary Figure 1.**
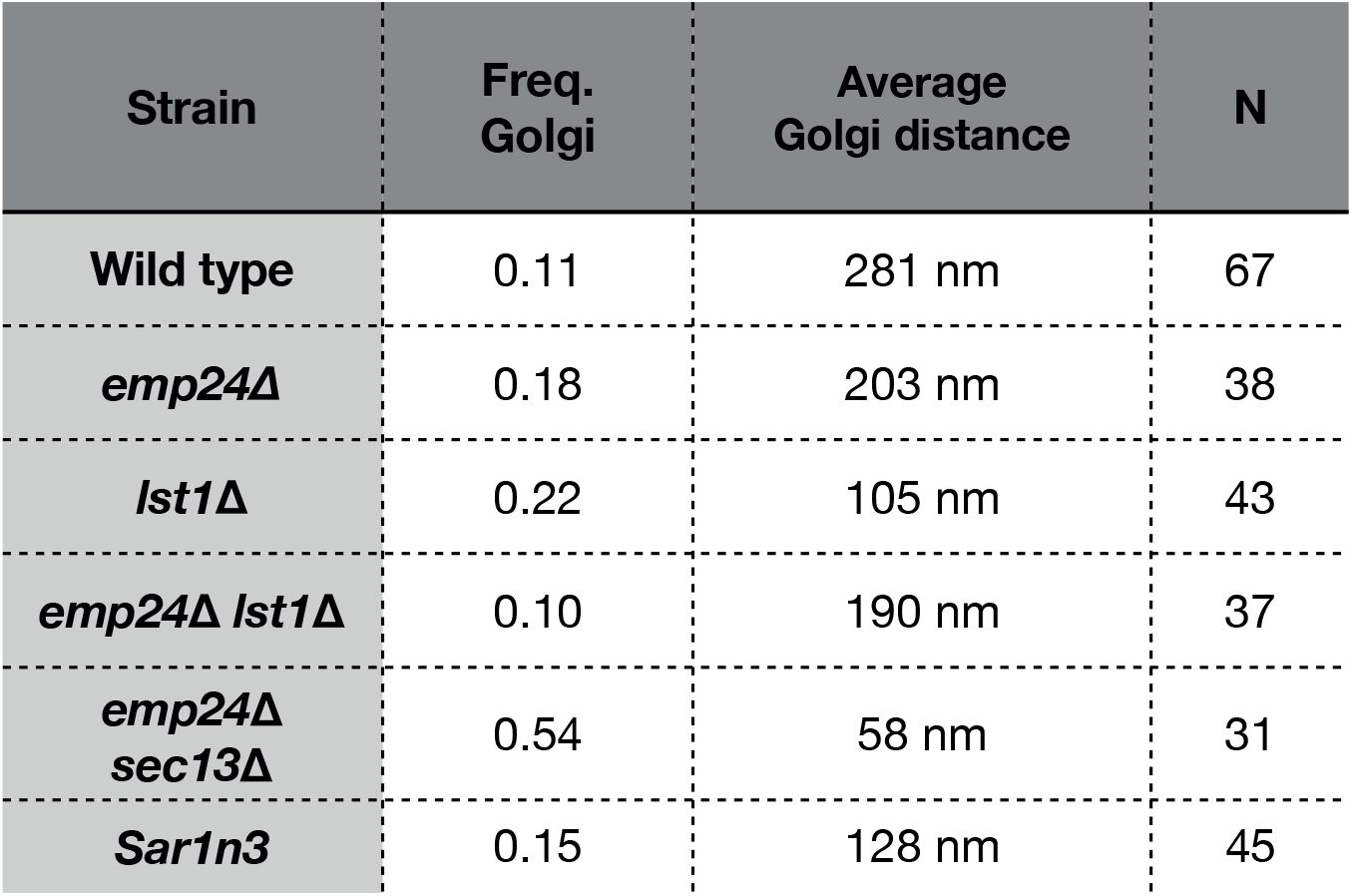
Table indicating the frequency and average distance to structures clearly identifiable as Golgi membranes. Frequency was calculated as the number of Sec16-sfGFP correlated spots with flattened Golgi complex cisterna within an unobstructed distance of <300nm (e.g., no other organelles obstructing the measurement). Distance was measured as the shortest distance from the Golgi cisterna to the relevant COPII structure or ER surface within 125nm of the correlated spot.

**Supplementary Figure 2.**
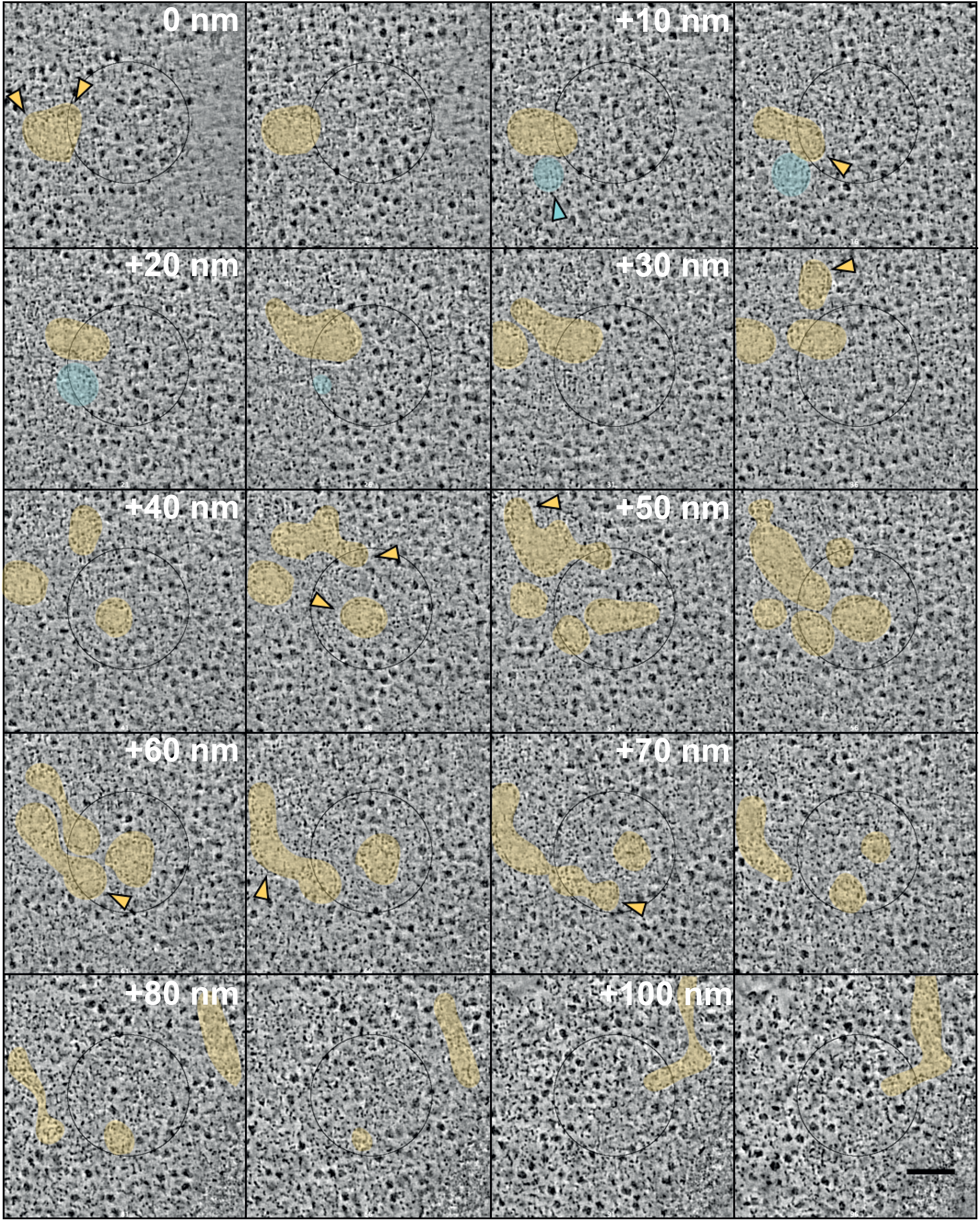
Composite of virtual slices from a tomogram from an *emp24*Δ *sec13*Δ *SEC16-sfGFP* strain. ER is highlighted in yellow, vesicles are highlighted in cyan, and arrowheads in corresponding colors point to buds and vesicles. The gap in z between each virtual slice is 5 nm. Scale bar is 100 nm.

## Notes

### Competing Interest Statement

The authors have declared no competing interest.

